# Histone deacetylation promotes transcriptional silencing at facultative heterochromatin

**DOI:** 10.1101/219535

**Authors:** Beth Rosina Watts, Sina Wittmann, Maxime Wery, Camille Gautier, Krzysztof Kus, Dong-Hyuk Heo, Cornelia Kilchert, Antonin Morillon, Lidia Vasiljeva

## Abstract

It is important to accurately regulate the expression of genes involved in development and environmental response. In the fission yeast *Schizosaccharomyces pombe*, meiotic genes are tightly repressed during vegetative growth. Despite being embedded in heterochromatin these genes are transcribed and believed to be repressed primarily at the level of RNA. However, the mechanism of facultative heterochromatin formation and the interplay with transcription regulation is not understood. We show genome-wide that HDAC-dependent histone deacetylation is a major determinant in transcriptional silencing of facultative heterochromatin domains. Indeed, mutation of class I/II HDACs leads to increased transcription of meiotic genes and accumulation of their mRNAs. Mechanistic dissection of the *pho1* gene where, in response to phosphate, transient facultative heterochromatin is established by overlapping lncRNA transcription shows that the Clr3 HDAC contributes to silencing independently of SHREC, but in an lncRNA-dependent manner. We propose that HDACs promote facultative heterochromatin by establishing alternative transcriptional silencing.

## Introduction

Heterochromatin is critical to eukaryotic cells and exists in two functionally distinct forms: constitutive and facultative heterochromatin. Constitutive heterochromatin is found mainly at gene-poor regions such as centromeres and telomeres, and is important for chromosome segregation during cell division. In contrast, facultative heterochromatin is found in gene-dense regions of chromosomes and controls silencing of developmentally and environmentally regulated genes (including meiotic genes^1–4^ and genes regulated by phosphate availability, such as *pho1*^5^). The mechanisms underlying the formation and maintenance of stable constitutive heterochromatin have been extensively studied in the fission yeast *Schizosaccharomyces pombe* (*S. pombe*), which has proven a powerful model for understanding heterochromatic silencing. In contrast, the mechanisms mediating silencing of reversible facultative heterochromatin are not well understood.

Chromatin modifying complexes play a central role in heterochromatin formation. These include the Clr4/Suv39h-methyltransferase complex (CLRC) that methylates Lys9 of histone H3 (H3K9me), as well as the class I (Clr6 and Hos2), class II (Clr3) and class III (Sir2, NAD^+^-dependent, ‘sirtuins’) histone deacetylase complexes (HDACs). Hypoacetylation of histones often depends on HDAC recruitment via low level non-coding (nc) transcription occurring within these regions. Heterochromatic ncRNAs are converted into dsRNAs through the action of the RNA-directed RNA polymerase complex (RDRC) and subsequently cleaved by the ribonuclease III Dicer (Dcr1) to produce small interfering RNAs (siRNAs) of 21-23 nucleotides^6–8^. These siRNAs are converted into single-stranded siRNAs and, together with Argonaut proteins, form the RNA-induced transcriptional silencing (RITS) complex. siRNAs direct the other RITS components, as well as Clr4, to heterochromatin by complementary base pairing with nascent transcripts^6,9,10^. The Clr4 H3K9me mark, in turn, acts as a landing platform for the HP1-like proteins Swi6 and Chp2^7^. These proteins target class I and II HDAC complexes, leading to further compaction of heterochromatin and transcriptional gene silencing^11^. The class I HDAC Clr6 removes acetyl groups from H3K14 and K9, and H4K5, K8, K12 and K16^12^. The class II HDAC Clr3 is part of SHREC (*S*nf2/*H*dac [histone deacetylase]-containing *Re*pressor *C*omplex) that deacetylates H3K14. In addition to Clr3, SHREC is composed of four other factors – Clr1, Clr2 and the ATPase and chromatin-remodelling subunit Mit1^11^. SHREC has been shown to be recruited to pericentromeric heterochromatin by the conserved protein Seb1^13^, which harbours RNA- and Pol II C-terminal-domain (CTD) binding modules (RNA-Recognition-Motif, RRM and CTD-Interacting-Domain, CID).

Unlike at constitutive heterochromatin, H3K9me at facultative heterochromatin does not induce transcriptional silencing. Instead, it has been proposed that meiotic transcripts are primarily silenced post-transcriptionally, with the RNA-binding proteins Mmi1 (which recognises DSR (Determinant of Selective Removal) sequences on RNA^14^) and Zinc-finger protein Red1 recruiting the RNA degradation machinery. Mmi1 and Red1 do also recruit RNAi and establish H3K9me but transcriptional silencing does not appear to be induced as a consequence. Despite all this, we previously demonstrated that the acid phosphatase-encoding gene *pho1*, on which we found facultative heterochromatin, is repressed at the transcriptional level by the DSR-containing lncRNA *prt* (*pho1-*repressing transcript). *prt* is produced in the presence of phosphate from a promoter located upstream of *pho1*^5,15–17^. Under such conditions, expression of *prt* leads to transcriptional repression of *pho1*. Co-transcriptional recruitment of Mmi1 to *prt* initiates the deposition of H3K9me by Clr4, formation of transient facultative heterochromatin across the gene and degradation of *prt* by the exosome complex. In the absence of non-coding transcription (either when the non-coding promoter is deleted or when cells are shifted to media without phosphate) *pho1* transcription is fully activated. Surprisingly though, in contrast to the complete loss of silencing observed upon deletion of the *prt* non-coding promoter, deletion of Clr4 had a minor effect on *pho1* expression and instead affected only induction kinetics^5^. This led us to speculate that there could be another, Clr4-independent mechanism of repression that is also mediated by *prt* non-coding transcription.

In *Saccharomyces cerevisiae* (*S. cerevisiae*) it has been shown that meiotic genes are repressed at the level of transcription by HDAC activity^18,19^. Repression of euchromatic genes by non-coding transcription has also recently been shown to require the class I HDAC Rpd3S (Rpd3 Small complex) and Set3/Rpd3L (Rpd3 Large complex)^20,21^. HDAC activity within these complexes is tightly linked to methylation of either H3K36 or H3K4 by, respectively, Set2 and Set1 methyltransferases. Through interacting with serine 5 phosphorylated CTD Set1 methylates H3K4 at gene 5’ ends^22^. It establishes a zone of H3K4me3 near transcription start sites followed by a zone of H3K4me2 downstream^23,24^. Set2 on the other hand establishes H3K36 methylation later during transcription and interacts with CTD phosphorylated at serines 2 and 5^25–27^. H3K36me can negatively affect transcription by targeting histone deacetylation by Rpd3S^28–30^. Both the Set2-Rpd3S and Set1-Rpd3L pathways have been shown to be important for modulating transcriptional dynamics during certain environmental changes in *S. cerevisiae*. Interestingly, while steady-state mRNA levels are not affected, gene induction occurs more rapidly upon carbon source shift in the absence of Set2. This is reminiscent of the situation in *S. pombe* where we also see faster *pho1* induction in a Clr4 mutant. Here, homologous HDAC activities are provided by Clr6 and Hos2 complexes but, unlike budding yeast Rpd3, Clr6 is essential for cell viability^31^. A possible reason for these differences might be due to the fact that Clr6 also functions in transcriptional silencing at constitutive heterochromatin together with Clr3 and the class III HDAC Sir2.

In this study we set out to investigate whether HDACs contribute to transcriptional silencing at regions of facultative heterochromatin. Using the *pho1* locus as a model of facultative heterochromatin, we demonstrate that transcription is repressed upon inhibition of class I/II HDACS using trichostatin A (TSA). This seems to be due to the action of Clr3 as its deletion has the same effect on *pho1* levels. Strikingly, Clr3-dependent repression of *pho1* is mediated via the lncRNA *prt*, which is needed for Clr3 recruitment to the loci. Interestingly, loss of Set1 and Set2 methyltransferases also leads to loss of transcriptional silencing, with the Set1 deletion having the most striking effect, suggesting that the role of Set1 in facilitating recruitment of HDACs via non-coding transcription is conserved in budding and fission yeast. In contrast to constitutive heterochromatin^13^, Clr3 is likely to mediate *pho1* silencing by a mechanism that acts in addition to Clr4. Strikingly, class I/II HDAC inhibition results in loss of transcriptional silencing of meiotic genes as analysed by NET-Seq and RNA-Seq. We conclude that HDACs have an important function in transcriptional silencing at facultative heterochromatin.

## Results

### The class II HDAC Clr3 contributes to *pho1* silencing

We previously reported that transcriptional repression of the acid phosphatase *pho1* in response to extracellular inorganic phosphate relies on the transcription of the overlapping ncRNA *prt*^5^. We showed that the H3K9 methyltransferase Clr4 contributes to this effect but, since its deletion does not lead to complete loss of silencing, we speculated that other players must be involved. In addition to H3K9me, hypoacetylation of histones by HDACs has been linked to transcriptional repression. To then investigate whether HDACs are involved in repression at *pho1*, we treated cells with a class I and II HDAC inhibitor, trichostatin A (TSA). Strikingly, a substantial increase in *pho1* mRNA is observed by Northern blot when cells are transiently treated with TSA (Figure 1a). At the same time, no change in the expression level of the house-keeping gene *adh1* is observed upon TSA treatment.

**Figure 1:**
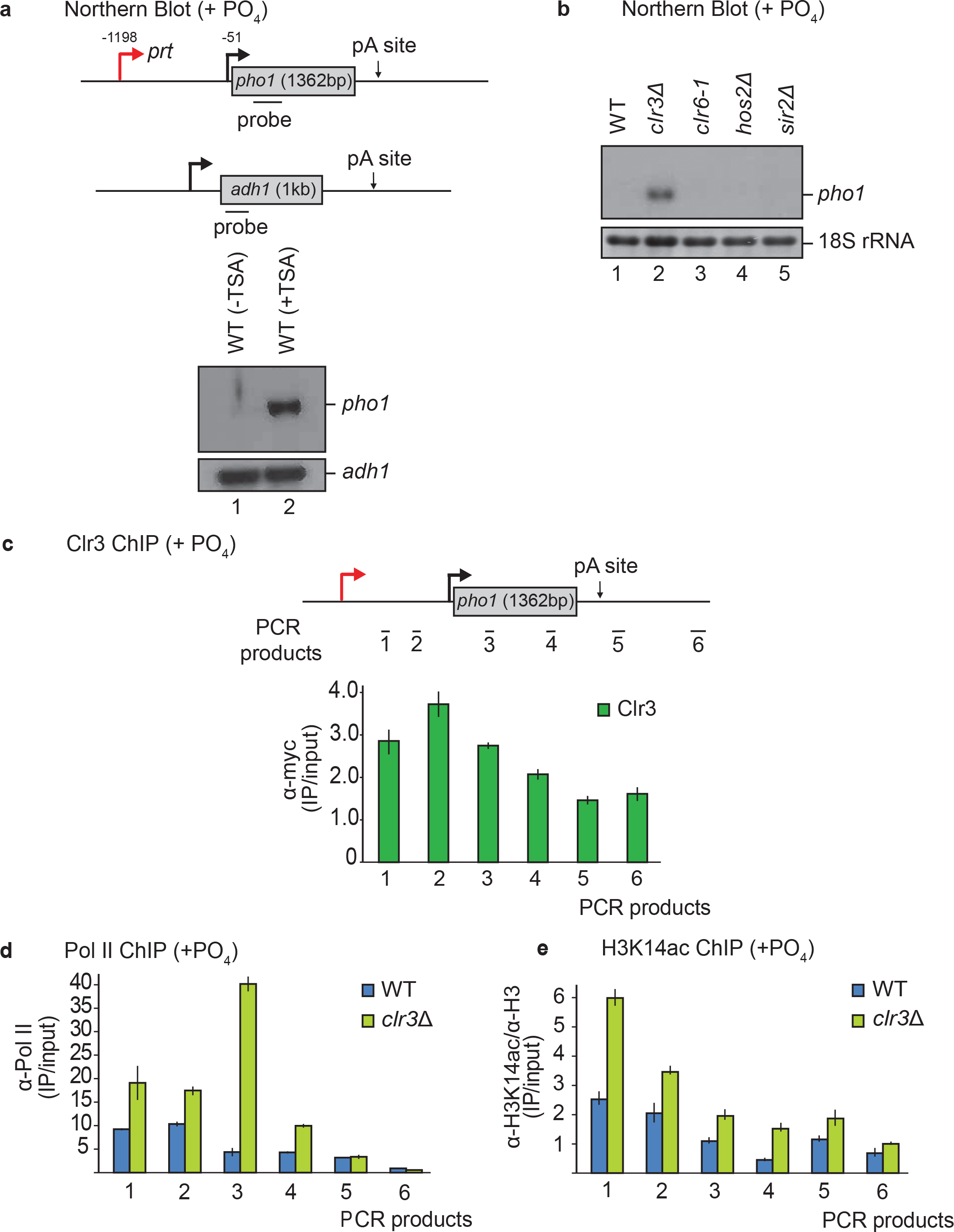
The HDAC Clr3 contributes to *pho1* silencing. (a) Schematic of *pho1* (top) and *adh1* (bottom) genes are shown with probes indicated by black bars. Northern blot analysis of RNA isolated from a wild-type (WT) strain untreated (-TSA) or treated (+TSA) with 20 μg/mL of the HDAC inhibitor trichostatin A (TSA). The housekeeping gene *adh1* serves as loading control. (b) Northern blot analysis of RNA isolated from the indicated strains grown in YES. 18S ribosomal RNA serves as a loading control. (c) Schematic of *pho1* gene with qPCR products indicated by black bars. Clr3-myc ChIP was performed in YES. (d) Pol II and (e) H3K14ac ChIP analysis performed in a *clr3*Δ strain compared to wild-type (WT) levels. In all ChIP-qPCR figures the mean from three independent experiments is shown; error bars indicate the standard error of the mean.

In order to identify which of the four *S. pombe* HDACs is responsible for the repression of *pho1*, we analysed RNA levels from the following HDAC mutant strains: *clr3*Δ, *clr6-1*, *hos2*Δ and *sir2*Δ (Figure 1b). Of the TSA-sensitive HDACs, Clr3 was the only one that had an effect on *pho1* expression, with its deletion resulting in increased *pho1* mRNA levels (Figure 1b). In contrast, deletion of TSA-insensitive *sir2*, which contributes to transcriptional silencing at constitutive heterochromatin, had no effect on *pho1* expression (Figure 1b, lane 5). Furthermore, no additional increase was observed when *clr3*Δ cells were treated with TSA suggesting that the increased *pho1* levels observed upon TSA treatment are solely due to Clr3 inhibition (Figure 3e, lanes 5 and 6). Previously, we showed that *pho1* levels are altered in RNA decay mutants (*rrp6*Δ and *mmi1*Δ) due to increased expression of *prt*^5^. In contrast, *clr3*Δ does not affect levels of *prt* (Figure 1b), suggesting a different mechanism of action here.

To further understand the role of Clr3 in repression of *pho1*, chromatin immunoprecipitation (ChIP) followed by qPCR was performed to determine whether Clr3 is recruited to the locus. This revealed that Clr3 indeed localizes to the gene, particularly at the non-coding region (Figure 1c). A recent study proposed a mechanism of Clr3 recruitment via the DNA and RNA binding activity of the non-catalytic SHREC subunit Clr2^32^. In order to determine whether Clr3 is recruited and functions at *pho1* as part of SHREC or independently of the complex, the effects of *clr1*, *clr2,* and *mit1* deletion on RNA levels were analysed by Northern blot (Figure Supplement 1a). Compared to wild-type *pho1* mRNA levels, a slight accumulation was detected for all strains. However, this is less pronounced than in *clr3*Δ, indicating that Clr3 is unlikely to entirely depend on Clr2 or other components of SHREC for recruitment to *pho1*. This is consistent with previous reports showing that Clr3 can also localize to several sites independently of other SHREC components^11^.

### Lack of Clr3 coincides with increased H3K14 acetylation and transcription at *pho1*

Having established that Clr3 is recruited to the non-coding *prt* region, we wanted to investigate how it exerts its silencing effect on *pho1*. Previous studies at the *mat* locus have proposed that Clr3 can restructure the chromatin environment in such a way as to restrict Pol II access^33^. In order to address if this is the case at the *pho1* region as well, the occupancy of Pol II at the locus was determined by ChIP-qPCR (Figure 1d). We found that loss of Clr3 leads to increased Pol II levels upstream of the *pho1* promoter and particularly across the gene body. This is consistent with increased transcription of *pho1* mRNA.

Since Clr3 possesses histone deacetylase activity, we expected changes in H3K14 acetylation (H3K14ac) levels at the *pho1* locus in *clr3*Δ. Indeed, an increase in the levels of H3K14ac could be detected by ChIP-qPCR (Figure 1e); similar to what has been shown previously at pericentromeric heterochromatin^11^. Concomitantly, the biggest increase in H3K14ac occurred over the non-coding region to which Clr3 is recruited. Taken together these data suggest that Clr3 functions in silencing the *pho1* mRNA by a mechanism that depends on its HDAC activity.

### Clr3 and Clr4 function independently to silence *pho1*

It has previously been suggested that Clr3, as part of SHREC, promotes H3K9me at pericentromeric heterochromatin and acts in the same pathway as the Clr4 methyltransferase^13^. Consequently, we next tested whether this is also the case at the *pho1* locus. In order to address this, we generated the double mutant *clr3*Δ*clr4*Δ. Remarkably, this mutant showed an additive accumulation of *pho1* mRNA levels (Figure 2a) and displayed a slow growth phenotype considerably more severe than either of the single mutants, *clr3*Δ or *clr4*Δ (Figure 2b). These data support the idea that Clr3 and Clr4 can act independently of each other to promote gene silencing at facultative heterochromatin, in contrast to the mechanism proposed at constitutive heterochromatin.

**Figure 2:**
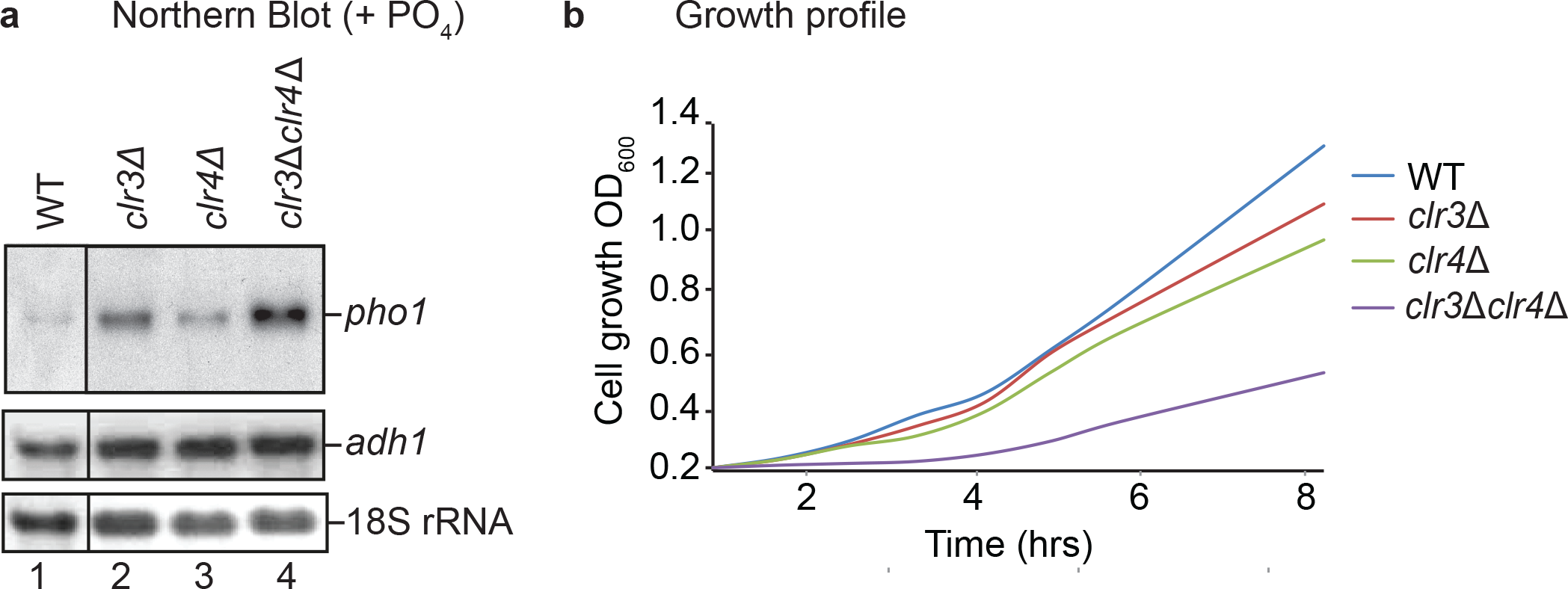
Cumulative effect of Clr3 and Clr4 on heterochromatin establishment. (a) Northern blot analysis of RNA isolated from the indicated strains grown in YES. *adh1* and 18S ribosomal RNA serve as loading controls. Probes used are as in Fig. 1a. (b) Growth profile of indicated strains grown in YES media over a period of eight hours.

### Non-coding transcription is required for Clr3 recruitment

To further our understanding of how Clr3 drives transcriptional silencing, we investigated whether non-coding transcription is needed for its function. We found that, in the absence of non-coding transcription in a strain lacking the *prt* promoter (*ncpro*Δ)^5^, Clr3 recruitment to *pho1* was reduced (Figure Supplement 2a). H3K14ac levels were also increased in this strain (Figure Supplement 2b), similar to that seen in *clr3*Δ (Figure 1e). Moreover, we reasoned that if Clr3 recruitment can only occur in the presence of non-coding transcription, a *clr3*Δ*ncpro*Δ double mutant should not have an additive effect on *pho1* expression compared to the respective single mutants. To test this, we analysed *pho1* levels in these strains and, as expected, Northern blot analysis revealed no obvious additive effect compared to the *ncpro*Δ single mutant (Figure Supplement 2c). This indicates that Clr3 is likely to operate downstream of *prt* transcription.

To test which specific region(s) of the *prt* ncRNA other than its Mmi1 binding site are involved in *pho1* silencing, a series of five different 189 bp deletions within the 5’ proximal region of *prt* were generated (*prt-1*Δ (-1197- to -1008), *prt-2*Δ (-1008- to -819), *prt-3*Δ (-819 to -630), *prt-4*Δ (-441 to -252), and *prt-5*Δ (-252 to -63) (*pho1* ATG=1)) (Figure 3a). We were unable to generate a deletion of the region -630 to -441. Interestingly, elevated *pho1* levels were observed in *prt-1*Δ, *prt-2*Δ, and *prt-3*Δ (Figure 3b, lanes 1-4) suggesting that element(s) responsible for *pho1* silencing are located within the 5’ part of *prt*. No change in *pho1* expression could be detected for either *prt-4*Δ or *prt-5*Δ (Figure 3b, lanes 5 and 6). In the case of *prt-3*Δ, this phenotype is likely a result of lost Mmi1 recruitment since this mutant lacks the DSR motifs we previously mapped (^5,34^, Figure 3a, Mmi1 CRAC). We tested Mmi1 recruitment to the *prt* locus in this mutant and it is indeed defective (data not shown). We next wanted to examine whether inhibition of Clr3 HDAC activity by TSA has any additive effect on *pho1* levels in the *prt* mutants. In *prt* mutants *prt-1*Δ, *prt-2*Δ, *prt-4*Δ, and *prt-5*Δ, *pho1* levels were identical to the wild-type upon TSA treatment (Figure 3b, lanes 7, 8, 9, 11, 12). In contrast, *pho1* levels in *prt-3*Δ (which lacks DSRs) were noticeably increased in TSA-treated compared to untreated cells (Figure 3b, compare lanes 4 and 10) or any other TSA treated samples. Taken together, these data are in agreement with a model in which Clr3 recruitment is independent of H3K9me and the Mmi1/Clr4/H3K9me pathway acts in parallel to Clr3/H3K14ac.

**Figure 3:**
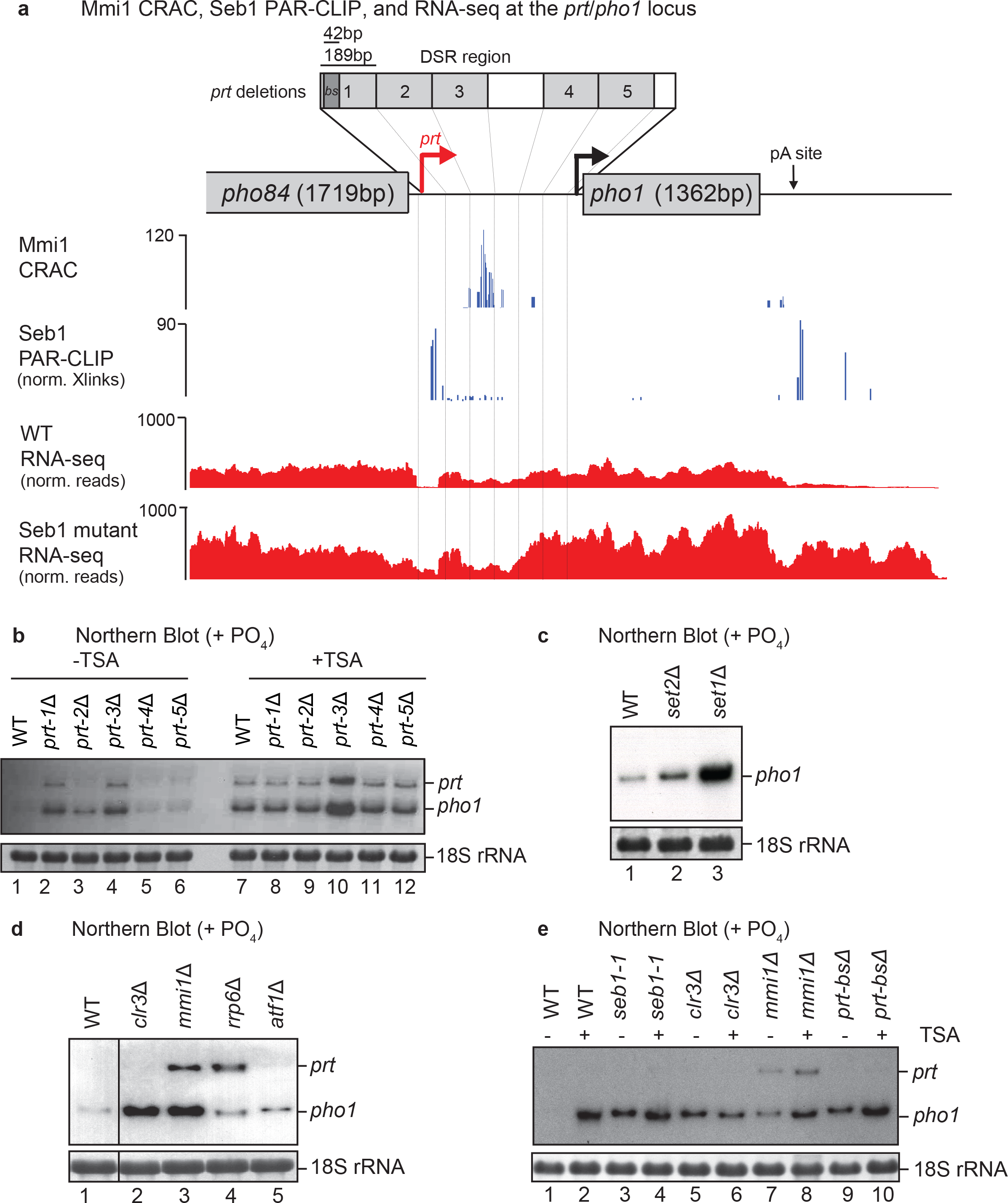
Mapping the contribution of specific regions of *prt* ncRNA to *pho1* silencing. (a) Schematic of *pho1* locus with deletions of *prt* regions as indicated by numbered grey boxes. Mmi1 CRAC and Seb1 PAR-CLIP signals are shown in blue and WT and Seb1 mutant S22D-K25E-K124E RNA-seq signals are shown in red. (b), (c), (d) and (e) Northern blot analysis of RNA isolated from the indicated strains grown in YES. 18S ribosomal RNA serves as a loading control. Probes used are the same as in Fig. 1a.

To understand further how transcription of *prt* mediates recruitment of Clr3 we wanted to test whether other histone methylation marks are involved in *pho1* repression. Interestingly, deletion of the H3K4 methyltransferase Set1 leads to derepression of *pho1* (Figure 3c). In contrast, deletion of the H3K36 methyltransferase Set2 only had a minor effect on *pho1* derepression, suggesting that the mechanism underlying *pho1* silencing primarily relies on histone methylation by Set1. In *S. cerevisiae*, Set1 mediates recruitment of the Rpd3L complex for repression of metabolic genes. However, we show that loss of the Rpd3 homologues Clr6 or Hos2 does not have an effect on *pho1* levels (Figure 1b). Instead, our data suggest that Set1 is likely to function via a different HDAC (Clr3) in transcriptional repression in fission yeast.

We next examined whether co-transcriptional recruitment of Clr3 is RNA dependent. To this end we performed ChIP experiments on chromatin extracts that were treated with RNases. As a control, we tested recruitment of the known *prt*-binding protein Mmi1 and, as expected, binding was lost upon RNase treatment (Figure Supplement 3a). We found the Clr3 signal at the *pho1* locus to be reduced suggesting that its recruitment is indeed dependent on RNA (Figure Supplement 3b). At the same time, RNase treatment did not affect the Pol II profile over *pho1* (Figure Supplement 3c). Interestingly, Set1 has recently been demonstrated to interact with nascent RNA within promoter–proximal regions via its RRM domains^35–37^. These studies proposed that RNA binding might be important for Set1’s function in transcriptional repression. It is possible that both Set1 and Clr3 interact with RNA. Alternatively, the observed loss of Clr3 from chromatin upon RNase treatment might be connected to loss of the methyltransferase, which is consistent with the model where HDAC targeting depends on Set1.

Given that budding yeast Set1 has no RNA sequence specificity, it is not entirely clear to us why deletion of the promoter-proximal regions of *prt* but not the distal regions leads to loss of *pho1* silencing (Figure 3b). Budding yeast Set1 was proposed to rely on Pol II phosphorylation (specifically on serine 5 within the CTD) for promoter-proximal targeting^38^. It is of course possible that the yeast proteins differ, and that *S. pombe* Set1 in fact has some preference either for nascent RNA sequence or secondary structure. Otherwise, specificity might be provided by another RNA or DNA-binding protein. Previous studies have shown that Clr3 recruitment can be mediated via the transcription factor Atf1^33,39,40^. However, the absence of the well characterized Atf1 binding site ATGACGT in *prt* suggests that Atf1 is unlikely to be involved in Clr3 recruitment to this locus. Indeed, no increase in *pho1* levels was observed in an *atf1*Δ strain (Figure 3d, lane 5). Altogether, these data suggest that HDAC-mediated silencing is dependent on Set1 and non-coding transcription and acts in addition to Mmi1/Clr4/H3K9me.

### HDACs act independently of RNA processing and degradation on *pho1*

To determine at which stage of RNA synthesis Clr3 mediates transcriptional silencing, we asked whether there is any genetic interaction between histone deacetylation and Seb1-dependent read-through or Mmi-dependent RNA degradation. We previously demonstrated that Seb1 is a key factor that regulates 3’ end cleavage and transcription termination of Pol II transcribed genes^41^. Accordingly, read-through transcription is observed in *seb1* mutants from the *pho84* gene located upstream of *prt* (^41^, Figure 3a), potentially interfering with its expression and leading to activation of *pho1*. Furthermore, consistent with Clr3 acting independently at this locus, an additive effect on *pho1* accumulation is observed when Clr3 HDAC activity is inhibited by TSA in strains either carrying mutation in *seb1* (*seb1-1*) or deletion of the Seb1 binding site (*prt-bs*Δ) (Figure 3e, compare lanes 3 to 4 and 9 to 10). Similarly, *mmi1*Δ shows an additive effect with TSA (Figure 3e, compare lane 7 to 8), which is also consistent with that seen when *prt-3*Δ, the strain harbouring a deletion spanning DSR elements (Figure 3a), is treated with TSA (Figure 3b, compare lanes 4 to 10). From these data we conclude that Clr3 is likely to act in addition to Seb1 and Mmi1 to repress *pho1*.

### Phosphate-response genes are regulated differently

To explore whether other genes involved in phosphate metabolism are regulated by HDACs, we studied the expression of a gene encoding *transporter for glycerophosphodiester 1* (*tgp1*). Similar to *pho1*, *tgp1* is repressed in response to inorganic phosphate, via non-coding transcription^16^, and induced upon growth in the absence of phosphate (Figure Supplement 4a, lanes 9-11). The *nc-tgp1* RNA is highly unstable and accumulates upon deletion of the exosome subunit Rrp6 (^16^, Figure Supplement 4a, lane 12) or the exosome specificity factor Mmi1^16,34^. We found accumulation of the *tgp1* transcript in the *seb1-1* mutant compared to wild-type (Figure Supplement 4a, compare lanes 1 and 2). In addition, we previously placed the *seb1* gene under the control of the thiamine regulated *nmt* promoter, where *seb1* expression is repressed upon switch from medium lacking to medium containing thiamine. Coincident with loss of Seb1, *tgp1* mRNA levels start to accrue (Figure Supplement 4a, lane 13). Seb1’s involvement in regulation of *tgp1* likely relates to failed termination of *nc-tgp1* in the two mutants, similar to the demonstrated role of its homologue in *S. cerevisiae*^42,43^. Unlike the case at *pho1,* we found that deletion of *clr3* has no discernible induction effect on *tgp1*. No detectable increase in either *tgp1* or *nc-tgp1* was observed in *sir2*Δ and two strains harbouring a *clr6* mutation (Figure Supplement 4a, lanes 6, 7 and 8) or upon treatment with TSA (Figure Supplement 4b). Thus, it appears that HDACs, or at least those sensitive to TSA, are not implicated in the repression of *tgp1* expression. Consistent with a *clr3* deletion strain having no effect on *tgp1* mRNA levels, recruitment of the protein, or indeed any other SHREC subunit, could not be detected at this locus (data not shown). Similarly, *clr4*Δ, or the double mutant *clr3*Δ*clr4*Δ (Figure Supplement 4a, lane 4 and 5), revealed no change in mRNA expression levels. These results indicate that unlike *pho1*, *tgp1* repression is not reliant on Clr3, other class I/II HDACs, nor Clr4's methyltransferase activity.

### Inhibition of HDACs induces dramatic changes in global Pol II transcription

We next asked how global expression of genes is regulated by the action of HDACs. To examine the contribution of class I and II HDACs on gene expression, we performed RNA-seq to systematically analyse the consequences of HDAC inhibition upon treatment with TSA. We used *S. cerevisiae* for spike-in to normalize for changes in overall RNA levels and compared cells before and after addition of TSA^44–46^. In contrast to our expectation considering the result at *pho1*, analysis of the data revealed that inhibition of HDAC activity results in more transcripts being decreased by at least 2-fold (3525) than increased (289) (Figure 4a and Figure Supplement 5a). Northern blot analysis confirmed decreased levels of *SPAC2E1P3.05c* and *ctr4* transcripts (Figure 4b) upon TSA treatment. A possible explanation for such a large amount of transcripts being decreased may relate to the role HDACs have been shown to have in suppressing overlapping antisense transcription in both fission and budding yeast^31,47^. Indeed, one study demonstrated that 74% of 23 transcripts positively regulated by the Rpd3L show overlapping antisense non-coding transcription^47^. However, our data did not show a strong correlation with antisense transcription for genes down-regulated upon inhibition of HDACs (data not shown). This suggests that suppression of antisense transcription via HDACs may not be a major mechanism to positively control transcription in fission yeast.

**Figure 4:**
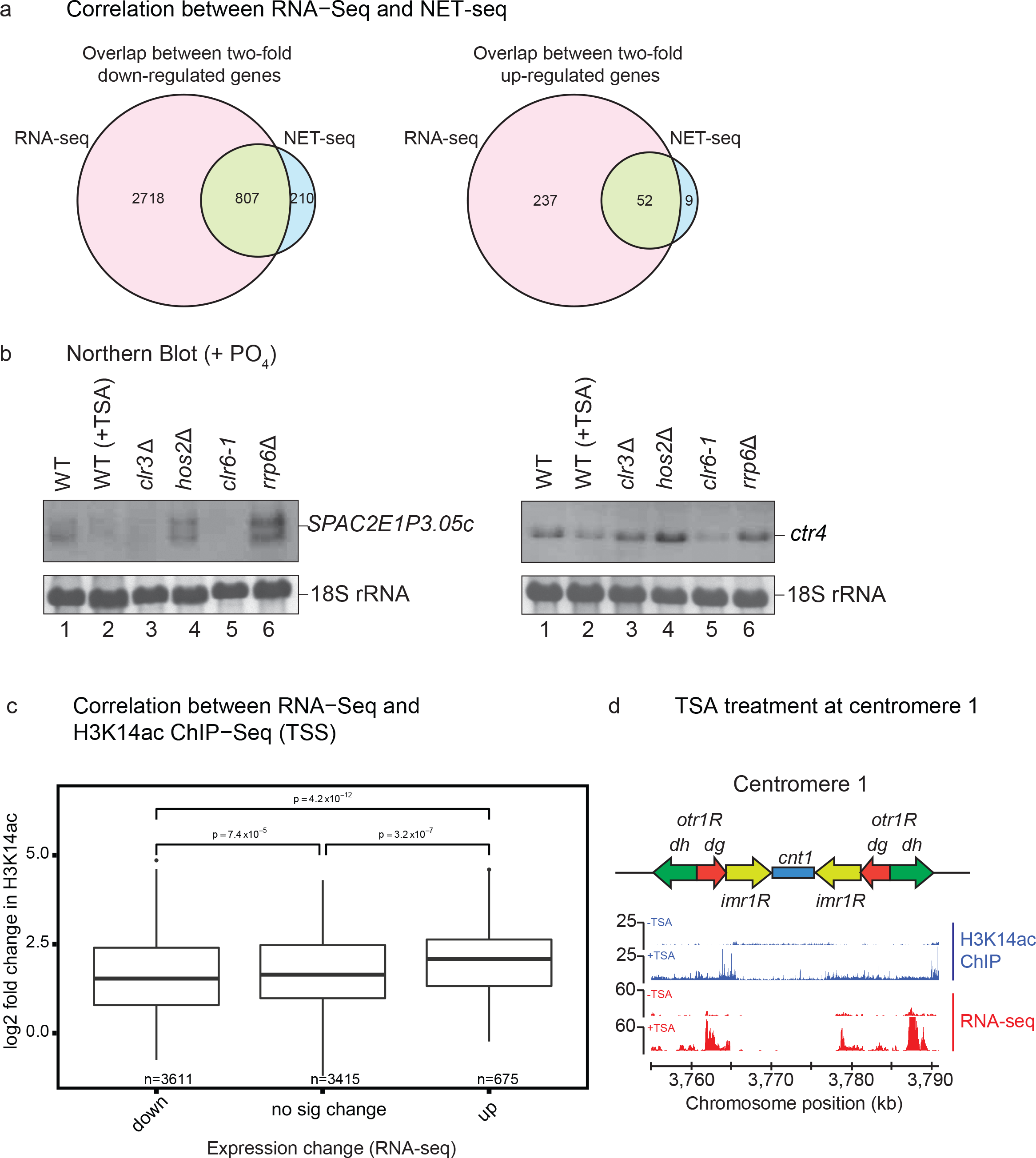
Genome-wide analyses. (a) Venn diagrams comparing genes identified by RNA-seq and NET-seq to be either down-regulated (left) or up-regulated (right) by treatment with TSA. (b) Northern blot analysis of RNA isolated from the indicated strains grown in YES and probed for *SPAC2E1P3.05c* (left) and *ctr4* (right). 18S ribosomal RNA serves as a loading control. (c) Genes were split, according to RNA-seq, into down-regulated, up-regulated or no significant change in TSA. The log2 fold change in H3K14ac of these different groups is shown as box plots. (d) Distribution of H3K14ac (blue) and RNA levels (red) at centromere 1 from untreated (-TSA) and treated (+TSA) samples. A map of *S. pombe* centromere 1 showing centromeric *otr1* regions comprising *dg* and *dh* repeats, *imr1* region and *cnt1* is shown.

In order to test whether the observed changes in transcript level are due to changed transcription or post-transcriptional RNA stability, as is the case in mammals^48^, we performed calibrated NET-seq. This technique allows for the genome-wide assessment of Pol II distribution at single-nucleotide resolution and in a strand specific manner^49^. We saw the same general trend of a greater number of genes being down-regulated than up-regulated (Figure Supplement 5a). Remarkably, about 80% of genes that are more than 2-fold up- or down-regulated show the same response to TSA in NET-Seq as observed by RNA-seq (Figure 4a). This suggests that the majority of changes in gene expression induced by HDAC inhibition occur at the transcriptional level.

### Increased histone acetylation correlates with transcription activation upon TSA treatment

To determine the direct effect of TSA treatment on gene expression and histone acetylation levels, we performed ChIP-seq experiments to measure histone H3K14ac and histone H3 levels genome-wide. While H3K14ac levels show large differences between TSA treated and untreated samples, the overall levels of H3 do not seem to be affected (Figure Supplement 5b). Consistent with previous results^50^ we identified a peak of H3K14ac at gene 5’ ends (150 nt before to 150 nt after gene TSS) (Figure Supplement 5c). Increased H3K14ac in this region correlates with up-regulation in RNA levels, which is in good agreement with a positive role for acetylation on gene transcription and a direct role for HDACs in repression of these genes (Figure 4c). Majority of the genome shows an increase in acetylation (Figure Supplement 5d). As expected this is true at centromeres and, as a consequence, we see higher RNA levels (Figure 4d). Consistently, down-regulated transcripts show lower H3K14ac levels compared to genes whose expression is not significantly affected by TSA treatment (Figure 4c). However, this effect is less important than the observed increase of H3K14ac for two-fold up-regulated genes, suggesting that lack of H3K14ac alone does not play a main role in transcription repression. It is possible that deacetylation at other lysines by Clr6 could be primarily responsible for a repressive effect on gene expression. Indeed, a strong decrease in the levels of the transcripts *SPAC2E1P3.05c* and *ctr4* can be seen upon mutation of Clr6 (Figure 4b). While deletion of Clr3 results in down-regulation of *SPAC2E1P3.05c* RNA this is not the case for *ctr4* suggesting that the observed effect upon TSA treatment is due to a different HDAC than Clr3. However, this does not seem to be Hos2 as no decrease in RNA levels is observed in a mutant of this HDAC.

### Transcriptional silencing of meiotic genes requires HDAC activity

Interestingly GO enrichment analysis of the genes up-regulated in TSA and with increased H3K14ac, revealed meiotic transcripts (Supplementary Table 1 and 2 and Figure 5a, b and c), and multiple membrane proteins including *efr3*, *pdh1*, *SPCC1235.18/17*, *SPAC977.06*, *ght5*, *ght6* and *ght8* (Supplementary Table 1 and 2 and Figure Supplement 6 and 7). Regions encoding for repetitive selfish elements also appear to depend on HDACs for transcriptional silencing. An interesting example is a group of so called selfish *wtf* genes (Figure Supplement 7). These proliferate due to meiotic drive, killing gametes that do not carry the genes and causing infertility^51^. Similar to *pho1* (Figure 5d), meiotic genes depend on Clr3 for transcriptional silencing. In the case of *meu31* some redundancy can be seen with Clr6 and Hos2 also contributing to the repression of transcription (Figure 5c). In agreement with a previously demonstrated role for the nuclear exosome complex in the degradation of meiotic transcripts during mitosis, we observe increased levels of *meu19* and *meu31* in *rrp6*Δ (Figure 5b and c). However, *mug14* is not affected in this exosome mutant, suggesting that the gene is likely to be regulated only at the transcriptional level (Figure 5a). In support of repression stemming from both the transcriptional and RNA level, serial dilutions onto TSA plates reveal a strong synthetic growth defect for *rrp6*Δ (Figure 5e). In summary, our data suggest that, in addition to post-transcriptional regulation, regions of facultative heterochromatin around *prt/pho1* and meiotic genes are repressed at the transcriptional level via the action of HDACs.

**Figure 5:**
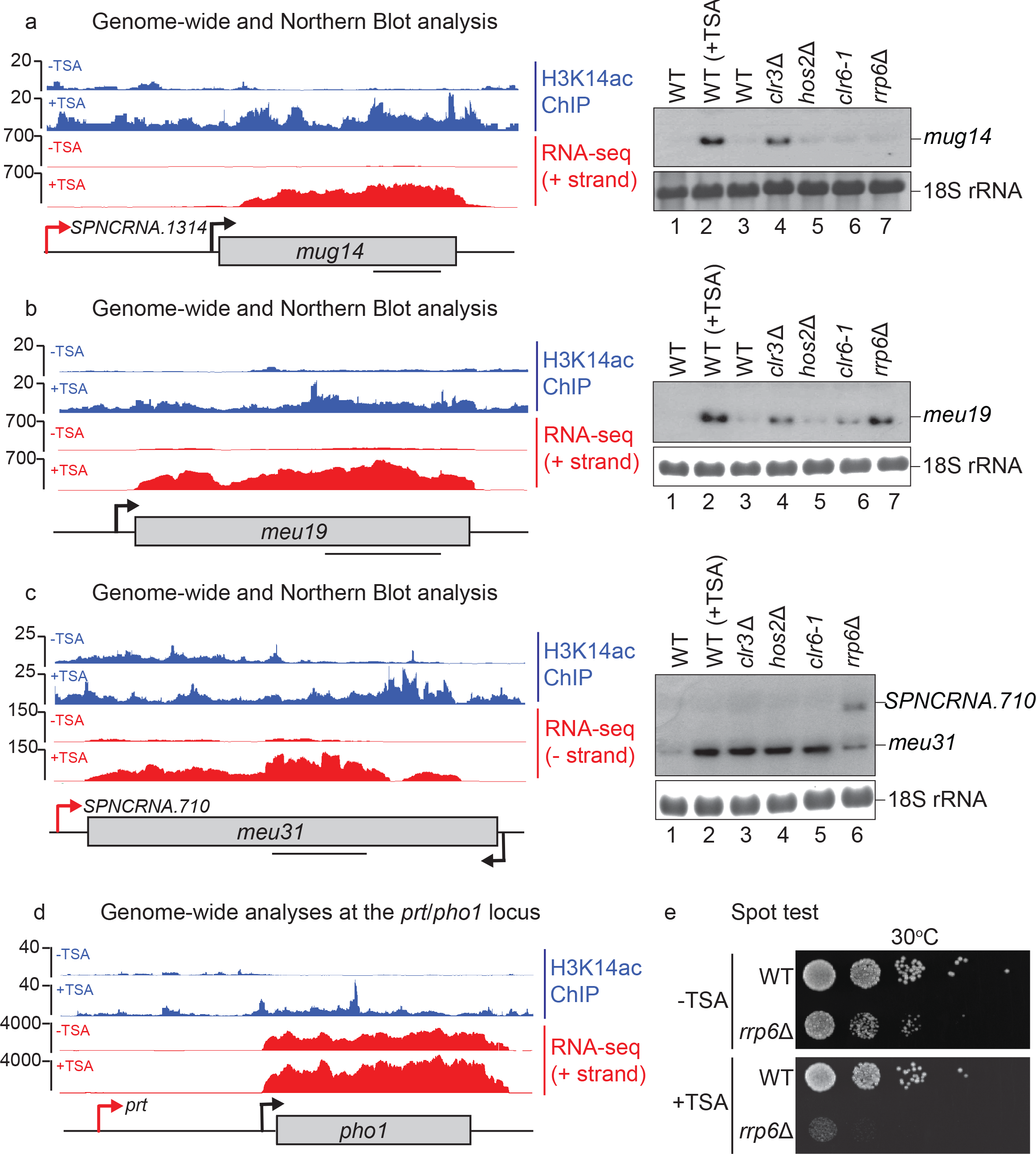
Transcriptional silencing of meiotic genes requires HDAC activity. Northern blot analysis (right) of RNA isolated from the indicated strains grown in YES and probed for (a) *mug14*, (b) *meu19*, and (c) *meu31*. 18S ribosomal RNA serves as a loading control. H3K14ac ChIP-seq (blue) and RNA-seq (red) are shown +/- TSA for each gene (left). Black bars on gene schematics indicate probes used for Northern blot. (d) Schematic of *prt/pho1* locus with H3K14ac ChIP-seq signal (blue) and RNA-seq reads (red) +/- TSA visualized using IGB. (e) Serial spot dilutions of WT and *rrp6*Δ strains spotted onto YES without TSA (-TSA) or with 20 μg/ml TSA (+TSA).

### *pho1*, an archetype of lncRNA/HDAC-mediated gene regulation?

Finally, we wanted to examine whether there are other loci that, similar to *pho1,* are controlled via unstable non-coding transcripts originating from intergenic regions. To first identify non-coding transcripts that are regulated by the exosome, we performed high resolution calibrated RNA-seq comparing wild-type with exosome deficient *rrp6*Δ. Our spike-in normalized results show up-regulation of multiple transcripts that were previously reported to be degraded by the exosome complex. We were also able to annotate new transcripts whose levels are controlled by the exosome (Supplementary Table 1). One such transcript is the novel intergenic cryptic unstable transcript (CUT) between the *SPAC27D7.09c/11c* loci (Figure Supplement 6). Interestingly, similar to meiotic genes, these CUTs are also up-regulated upon inhibition of HDAC activity. Interestingly, at least for some of the transcripts that overlap with ncRNA, we observe increases in H3K14 acetylation and Pol II transcription (Supplementary Table 3 and Figure Supplement 6 and 7). These transcripts include genes encoding for membrane transporters, such as the calcium transporter *cta3*, and hexose transporters, such as *ght8.* This suggests that other loci might be regulated via HDAC activity dependent on non-coding transcription, possibly via the activity of the Rpd3 (Clr6 and Hos2) complexes described in *S. cerevisiae*.

We conclude that class I and II HDACs play a significant role in shaping the fission yeast transcriptome by regulating transcription both positively and negatively. We have demonstrated that regions of facultative heterochromatin, including meiotic genes and genes regulated via non-coding transcription, such as *pho1*, require HDAC activity for their transcriptional repression.

## Discussion

The ability of cells to invoke rapid changes in gene expression is critical for the execution of developmental programs, adaptation to environmental stress, and during the fate decision of stem cells^52^. Many meiotic mRNAs are repressed in mitotic cells and derepressed during meiosis when the function of the proteins they encode is needed. Similarly, expression of the genes encoding for metabolic enzymes such as *pho1* is also dynamically regulated in response to nutrient availability^5^. Despite these genes being rooted in a chromatin environment that has the canonical features of heterochromatin (such as H3K9me and marks associated with transcriptional repression at centromeric and telomeric regions), loss of H3K9me upon deletion of Clr4 does not lead to up-regulation of their transcription^4,5,16^. Instead, it was believed that they were repressed primarily at the level of RNA stability. Here, we demonstrate that *pho1* and meiotic genes are in fact repressed at the level of transcription by HDACs. This repression works in concert with RNA degradation and H3K9me (Figure 6). Indeed, we see a synthetic growth defect when suppression of HDACs (TSA treatment) is combined with inhibition of RNA degradation (*rrp6*Δ).

How HDACs control transcriptional repression of only specific regions of the genome is not fully understood. Recent studies have proposed that the pattern of histone methylation at promoters of protein-coding genes could provide the specificity. Histone H3 methylation by Set1 at promoter proximal regions controls gene expression either positively or negatively depending on the H3K4me2/me3 ratio^23,24^. A characteristic chromatin environment featuring high levels of H3K4me2 has been proposed to recruit the Rpd3L HDAC to repress transcription in budding yeast^20^, whereas H3K4me3 positively affects transcription. H3K4me2 peaks downstream of H3K4me3 and represses initiation at cryptic promoters to promote the production of full-length mRNA^47^. According to this model, unless H3K4me2 is introduced through the act of upstream overlapping non-coding transcription, genuine promoters should be resistant to Set1-dependent inhibition. We demonstrate that transcriptional silencing within facultative heterochromatin, in fission yeast, is established via non-coding transcription and relies on a Set1/Clr3 mechanism. Multiple non-coding transcripts recently reported to be produced from regions proximal to or overlapping meiotic transcripts^53–55^ might be responsible for mediating the effect of Set1 and Set2 in recruitment of class I/II HDACs to these genes. However, it is not entirely clear how widespread the role of non-coding transcription is in repressing regions of facultative heterochromatin derepressed by TSA. We show examples of protein-coding and non-coding transcripts that do not have any overlapping non-coding transcription and whose promoters are likely to be directly regulated by Set1. While it is not clear how Set1 represses genuine promoters one possibility could be that the ratio of H3K4me2/me3 is higher on these genes due to different Set1 residency times. Indeed, recent studies have shown that residency time on promoters can be regulated via Set1’s interaction with the nascent RNA. It was proposed that this could, in turn, affect the H3K4me2/me3 ratio^36,37^. This suggests an exciting possibility that Set1 binding to RNA is regulated at these promoters to create a specific chromatin environment that is not permissive for transcription. Alternatively, H3K4me2/me3 ratio could be regulated via demethylation activities targeted to these promoters.

Pervasive antisense transcription occurring as a result of Set2 deletion has been shown to modulate expression of highly regulated metabolic genes^56^, even though global antisense transcription does not show correlation with sense transcription^57^. Set2 deletion has also been shown to contribute to the regulation of *tgp1* and *pho1*^16^, although we show that Set2’s contribution at *pho1* is minor compared to Set1’s. We have shown that Rpd3 complexes and other class I/II HDACs are not involved in *tgp1* repression, suggesting that Set2 may repress this gene through a different mechanism. In contrast to constitutive heterochromatin where HDAC recruitment is dependent on H3K9me and sequence-specific DNA binding proteins (such as ATF/CREB family transcription factors Atf1 and Pcr1^39^), at facultative heterochromatin Clr4-dependent deposition of H3K9me depends on the sequence-specific RNA-binding protein Mmi1^34^. It has been proposed that Set1 can also be recruited to select euchromatic loci via Atf1^40^. While our data suggest that the *prt* promoter proximal region is important for *pho1* repression, we did not observe any effect on *pho1* in an *atf1*Δ strain, arguing against a role for Atf1 in providing the specificity for recruitment of Set1. Furthermore, transcripts whose expression was reported to be increased upon deletion of Atf1 differ from the transcripts we report to be derepressed upon HDAC inhibition.

It has previously been proposed that SHREC promotes H3K9me^13^. The mechanistic details of how SHREC achieves this are unclear but it has been suggested that either deacetylation of H3 or of a non-histone substrate by Clr3 is necessary for Clr4-mediated methylation of H3K9. In contrast to the case at constitutive heterochromatin, we show that Clr3 and Clr4 proteins act in parallel, and that H3K14 deacetylation plays a much more prominent role than H3K9 methylation in transcriptional repression. In mammals, developmental genes are believed to be primarily silenced by H3K27me3 positioned by Polycomb Repressive Complex 2 (PRC2) in proliferating cells. However, this has been challenged by a recent study demonstrating that H3K27me-independent histone deacetylation plays an important role in transcriptional repression of developmental genes in mouse cardiomyocytes, and is important for normal heart function^58^. This implies that the role of HDACs in transcriptional silencing of developmentally regulated genes is likely conserved in mammals and therefore it will be even more important to understand how these complexes function.

This study significantly advances our understanding of the molecular mechanisms underlying repression of meiotic genes. However, many questions are still left unanswered. It remains to be seen how HDACs are targeted to regions of facultative heterochromatin and why it is that many genes are down-regulated upon loss of HDAC activity. Future studies addressing some of these unresolved questions will provide further insight into the function of HDACs in the regulation of transcription.

## Materials and methods

### Yeast strains and manipulations

*S. pombe* strains were grown in either YES medium, or EMMG with 10 mM KH_2_PO_4_ to an OD_600_ of 0.4 - 0.7 before harvesting^59^. Strains and oligonucleotides used in this study are listed in Supplementary Tables 4 and 5.

Standard PCR-based methodology was used for epitope tagging^60^. The pop-in, pop-out method for allele replacement was used to introduce deletions of the *pho1* lncRNA^61^. DNA fragments carrying the desired deletions were generated by 2-step PCR. Fragments were confirmed by sequencing, and cloned into pCR blunt II TOPO (Life Technologies), according to the manufacturer’s instructions, before subcloning into pKS-URA4^60^ by SpeI/NotI digestion and ligation. BseRI was used to linearize the plasmid before transformation into *S. pombe* cells lacking the *ura4* locus. Ura^+^ colonies were selected on EMMG without uracil and genotyped by DNA sequence analysis of PCR products. Positive clones were plated onto 5-FOA to select cells in which the *ura4^+^* gene had been ‘popped-out’. Clones were verified by colony PCR and sequencing of products.

### Northern blotting

Northern blot experiments were essentially performed as described^62^. Gene-specific PCR-generated fragments were used as probes using oligonucleotides listed in Supplementary Table 5.

### Chromatin immunoprecipitation (ChIP)

ChIP was performed as previously described^5^. Immunoprecipitations (IPs) were conducted with either rabbit IgG agarose (Sigma) or antibodies against H3 (Abcam, 1791, RRID:AB_302613), H3K14ac (Millipore, 07-353, RRID:AB_310545), Rpb1 (Millipore, 8WG16, RRID:AB_492629) or c-Myc (Santa Cruz, SC-40, RRID:AB_627268) coupled to protein G dynabeads (Life Technologies). Primers for qPCR analysis are listed in Supplementary Table 5.

### RNase ChIP

RNase ChIP was performed as described above with the following exceptions: (i) formaldehyde cross-linking time was reduced from 20 min to 5 min (ii) Chromatin was prepared in FA lysis buffer with 0.05% SDS and (iii) cross-linked chromatin was treated with either 7.5 U of RNase A (Thermo Scientific) and 300 U of RNase T1 (Thermo Scientific) or an equivalent volume of RNase storage buffer (50 mM Tris-HCl (pH 7.4) and 50% (v/v) glycerol). After incubation at room temperature for 30 min, immunoprecipitations were performed as usual.

### ChIP-seq

Chromatin was prepared as above. After washing and eluting bound material from the beads, protein was removed by incubation with 0.4 mg pronase for 1 hr at 42°C, followed by overnight incubation at 65°C. RNA was degraded by incubating samples with 0.02 mg RNase A (Roche) for 1 hr at 37°C. DNA was purified using ChIP DNA Clean & Concentrator kit (Zymo Research, USA) according to the manufacturer’s instructions. A sequencing library was constructed using NEBNext Fast DNA Library Prep Set for Ion Torrent^TM^ Kit (NEB, USA). Libraries with different barcodes were pooled together and loaded onto the Ion PI^TM^ Chip v3 using the Ion Chef^TM^ Instrument (Life Technologies, USA). Library sequencing was carried out on the Ion Torrent Proton. The resulting sequences were trimmed to remove low quality reads (less than Phred score 20) and reads shorter than 20 nt using Trimmomatic (version 0.36)^63^.

### RNA sequencing

RNA-seq was performed in duplicates as described previously^34^. Libraries were prepared and sequenced by the High-Throughput Genomics Group at the Wellcome Trust Centre for Human Genetics on the Illumina HiSeq 2500 platform. Quality trimming was performed using Trimmomatic (Galaxy Version 0.32.3, RRID:SCR_011848)^63^.

For identification of CUTs, total RNA was extracted from biological duplicates of WT and *rrp6*Δ cells, using standard hot-phenol procedure. ERCC RNA Spike-In (2 μl of 1:100 dilution; Ambion) was added to 1 μg of total RNA, then rRNAs were depleted using the RiboMinus Eukaryote System v2 (Life Technologies). Strand-specific total RNA-Seq libraries were prepared using the TruSeq Stranded Total RNA Sample Preparation Kit (Illumina). Paired-end sequencing (2x50 nt) of the libraries was performed on a HiSeq 2500 sequencer.

### NET-seq

Biological duplicates of Rpb3-Flag *S. pombe* cells were treated for 2.5 hours with 20 μg/ml of the HDAC inhibitor trichostatin A (TSA). After lysis, each *S. pombe* lysate was mixed with an aliquot of a lysate of Rpb3-Flag *S. cerevisiae* cells (spike), in a 50:1 ratio.

Libraries were constructed according to the previously published protocol^49^, with minor modifications, starting from 1L of exponentially growing cells. Ligation of DNA 3’-linker was performed as described^64^, starting with 2-3 μg of purified nascent RNA. Ligated nascent RNA was then submitted to alkaline fragmentation for 20 minutes at 95°C. Single-end sequencing (50 nt) of the libraries was performed on a HiSeq 2500 sequencer.

After removal of the 5’-adapter sequence using cutadapt, reads were uniquely mapped to the *S. pombe* and *S. cerevisiae* reference genomes using version 2.2.5 of Bowtie, with a tolerance of 1 mismatch. Bioinformatics analyses used uniquely mapped reads. Tags densities were normalized either on the total number of uniquely mapped reads (IP & Input samples), or the signal for *S. cerevisiae* ribosomal proteins-coding genes.

### Processing and bioinformatic analysis of genome-wide data

Reads were aligned to the *S. pombe* genome (ASM294v2.28) and the *S. cerevisiae* genome (sacCer3) separately either with TopHat (Galaxy Version 0.9, RRID:SCR_013035)^65^ in the case of RNA-Seq, or with Bowtie2 v2.2.6 (RRID:SCR_005476)^66^ in the cases of NET-Seq and ChIP-Seq. Reads that were present in both alignments were removed using the Picard Tool Suite (http://broadinstitute.github.io/picard) to keep only reads unique for one of the genomes. All remaining *S. cerevisiae* reads were counted and used for the normalization of *S. pombe* samples. In the case of ChIP-Seq, differences in the input was also taken into account when calculating normalization values and H3K14ac was normalized to H3 density as determined by H3 ChIP-Seq. For NET-Seq, reads were trimmed to leave only the 3’ nucleotide corresponding to the nucleotide in the active centre of Pol II at the time of the experiment. For RNA-Seq, only properly aligned pairs were kept and all other reads were removed using samtools^67^. Spearman correlation matrices and metagene plots were calculated using deepTools^68^. For differential expression analysis, indicated regions were counted using R for ChIP-Seq and NET-Seq while HTSeq was used for RNA-Seq data^69^. The analysis for all data sets was performed with the R package DESeq2 (RRID:SCR_000154)^70^ using manual normalization to *S. cerevisiae* spike-in counts determined as described above. All other data analysis was also performed with R using in-house scripts featuring Bioconductor packages^71,72^.

For CUTs annotation, reads were mapped to the *S. pombe* reference genome using version 2.0.6 of TopHat, with a tolerance of 3 mismatches and a maximum size for introns of 5 kb. Tags densities were normalized on the ERCC Spike-In signals. Segmentation was performed using the ZINAR algorithm^73^. CUTs were defined as ≥200 nt segments showing a >2-fold enrichment in *rrp6*Δ vs WT, with a *P*-value (adjusted for multiple testing with the Benjamini-Hochberg procedure) <0.05 upon differential expression analysis using DESeq^74^.

### Data access

Raw (fastq) and processed sequencing data can be downloaded from the NCBI Gene Expression Omnibus repository (http://www.ncbi.nlm.nih.gov/geo, accession number GSE104713 (RNA-seq), GSE104712 (NET-seq).

## Acknowledgements

We thank H.D Madhani, and the National BioResource Project (NBRP) for strains. We are grateful to Sofia Battaglia and Patrick Cramer for contributions not shown here. This work was supported by the Wellcome Trust Research and Career Development and Wellcome Trust Senior Research fellowships to L.V. (WT088359MA and WT106994MA). High throughput sequencing was performed by the High-Throughput Genomics Group at the Wellcome Trust Centre for Human Genetics (Wellcome Trust grant 090532/Z/09/Z). This work has benefited from the facilities and expertise of the High-Throughput Sequencing platform of Institut Curie (Paris), supported by the Agence Nationale de la Recherche (ANR-10-EQPX-03, ANR10-INBS-09-08). The BBSRC provided support to B.R.W. and S.W. was supported by a studentship from the MRC.

## Contributions

B.R.W. and L.V. conceived and designed experiments. B.R.W. performed all experiments except *rrp6*Δ RNA-seq which was performed by M.W. and analysed by C.G. S.W. analysed ChIP-seq, RNA-seq, PAR-CLIP and NET-seq datasets. K.K and D-H.H also contributed to genome-wide analyses. B.R.W. and L.V. wrote the paper and all authors edited the manuscript.

## Supplementary File 1

**Supplementary Table 1:** RNA-, NET-, and ChIP-seq datasets

**Supplementary Table 2:** GO analysis of genes with increased transcription and H3K14ac

**Supplementary Table 3:** List of CUTs overlapping ORFs

**Supplementary Table 4:** List of *S. pombe* strains used in this study.

**Supplementary Table 5:** List of *S. pombe* oligonucleotides used in this study.

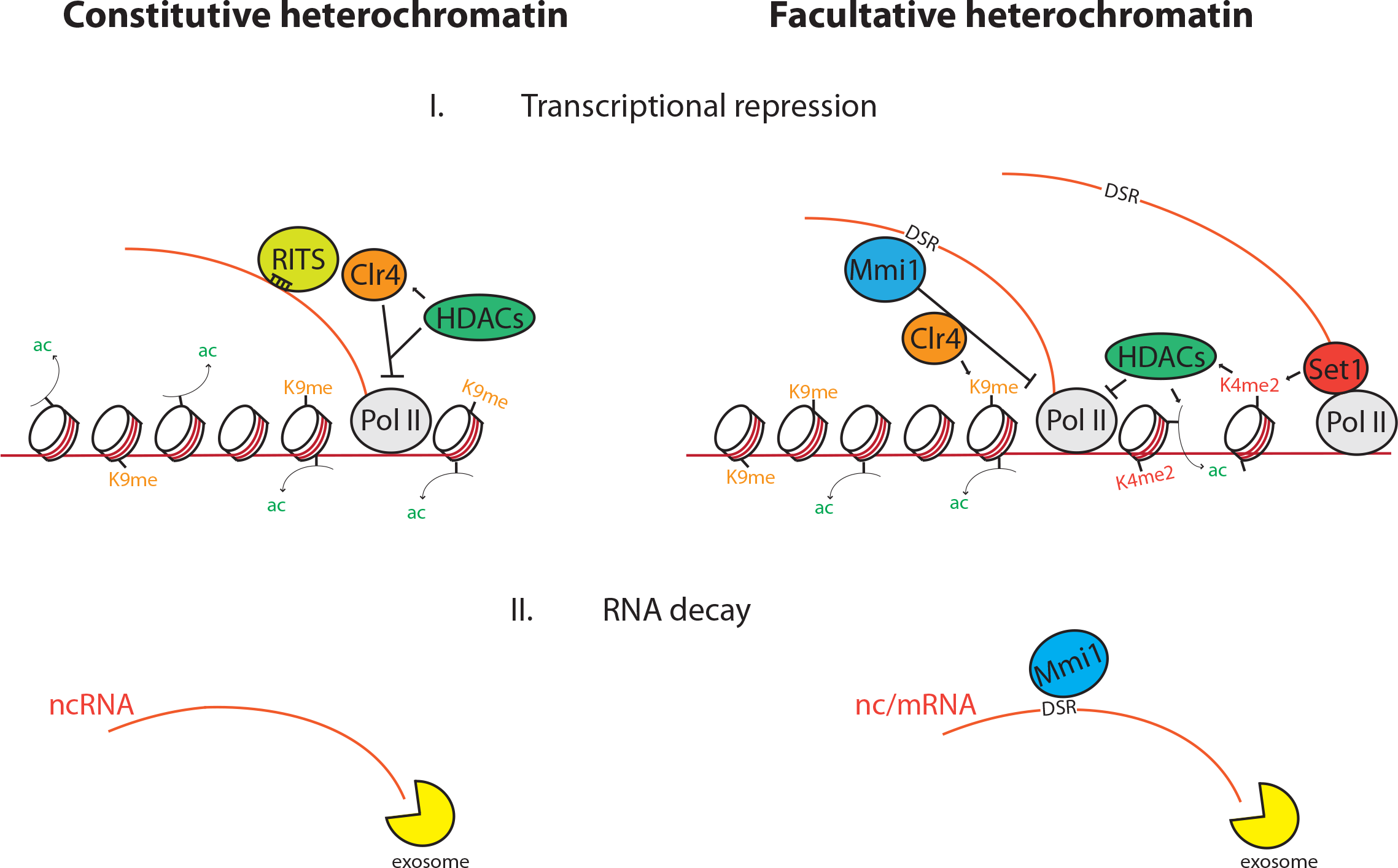

